# SLO potassium channels antagonize premature decision making in *C. elegans*

**DOI:** 10.1101/243220

**Authors:** Ichiro Aoki, Michihiro Tateyama, Takushi Shimomura, Kunio lhara, Yoshihiro Kubo, Shunji Nakano, Ikue Mori

## Abstract

**Summary:** Animals have to modify their behavior at the right timing to respond to changes in environments. Yet, the molecular and neural mechanisms regulating the timing of behavioral transition remain largely unknown. Performing forward genetics on a plasticity of thermotaxis behavior in *C. elegans*, we demonstrated that SLO potassium channels together with a cyclic nucleotide-gated channel CNG-3 determine the timing of the transition of temperature preference after shift of cultivation temperature. We further revealed that SLO and CNG-3 channels regulate the alteration in responsiveness of thermosensory neurons. Our results suggest that the regulation of sensory adaptation is a major determinant of the latency for animals to make decisions in changing behavior.

**Highlights:**

- Slo-1 and SLO-2 K^+^ channels decelerated transition of temperature preference in thermotaxis behavior after upshift of cultivation temperature
- SLO K^+^ channels slowed down the adaptation of AFD thermosensory neuron to new cultivation temperature
- A cyclic nucleotide-gated channel CNG-3 functioned together with SLO-2
- Thermotaxis serves as could be a model system for early onset epilepsies

## Introduction

One of the central issues in decision making is what determines the timing of behavior transition at which animals learn from a new experience and abandon the old knowledge (Doya, 2008; Wang, 2014). Appropriate latency for behavior transition is necessary for animals to discriminate a long-lasting environmental change from a transient noise, for instance, when animals emerge from hibernation (Williams et al., 2014). Conversely, if the latency is too long, animals would lose opportunities to obtain rewards or fail to avoid dangers. Although the regulation of the timing of behavior transition in response to environmental change is critical for animals, underlying molecular and neural principles remain largely unknown.

To understand how the timing of behavior transition is determined, we focused on a plasticity of thermotaxis behavior of *C. elegans*, in which animals associating cultivation temperature with the existence of food migrate toward that temperature on a thermal gradient with no food (Hedgecock and Russell, 1975; Mori and Ohshima, 1995, Figure 1A). Thermotaxis of *C. elegans* is essentially regulated by a simple neural circuit (Mori and Ohshima, 1995; Kuhara et al., 2008, Figure 1B). In this circuit, AFD is a major thermosensory neuron, which increases intracellular Ca^2+^ concentration ([Ca^2+^]_i_) in response to a temperature rise above the past cultivation temperature (Kimura et al., 2004; Kobayashi et al., 2016; Ramot et al., 2008; Clark et al., 2006; Wasserman et al., 2011). Since this cultivation temperature-dependency in AFD dynamic range is well conserved even when AFD cells isolated from the neural network is cultured *in vitro*, AFD encodes the information of cultivation temperature cell-autonomously (Kobayashi et al., 2016). Genetic analyses have revealed molecular components involved in temperature sensation in AFD. Three receptor-type guanylyl cyclases (GCs), GCY-8, GCY-18 and GCY-23, are specifically localized to the sensory ending of AFD and are thought to act as thermosensors (Inada et al., 2006; Takeishi et al., 2016). These GCs synthesize cGMP, which activates cyclic nucleotide-gated (CNG) channels composed of TAX-2 and TAX-4 (Komatsu et al., 1996; Coburn and Bargmann, 1996), leading to the Ca^2+^ influx into AFD.

**Figure 1.**
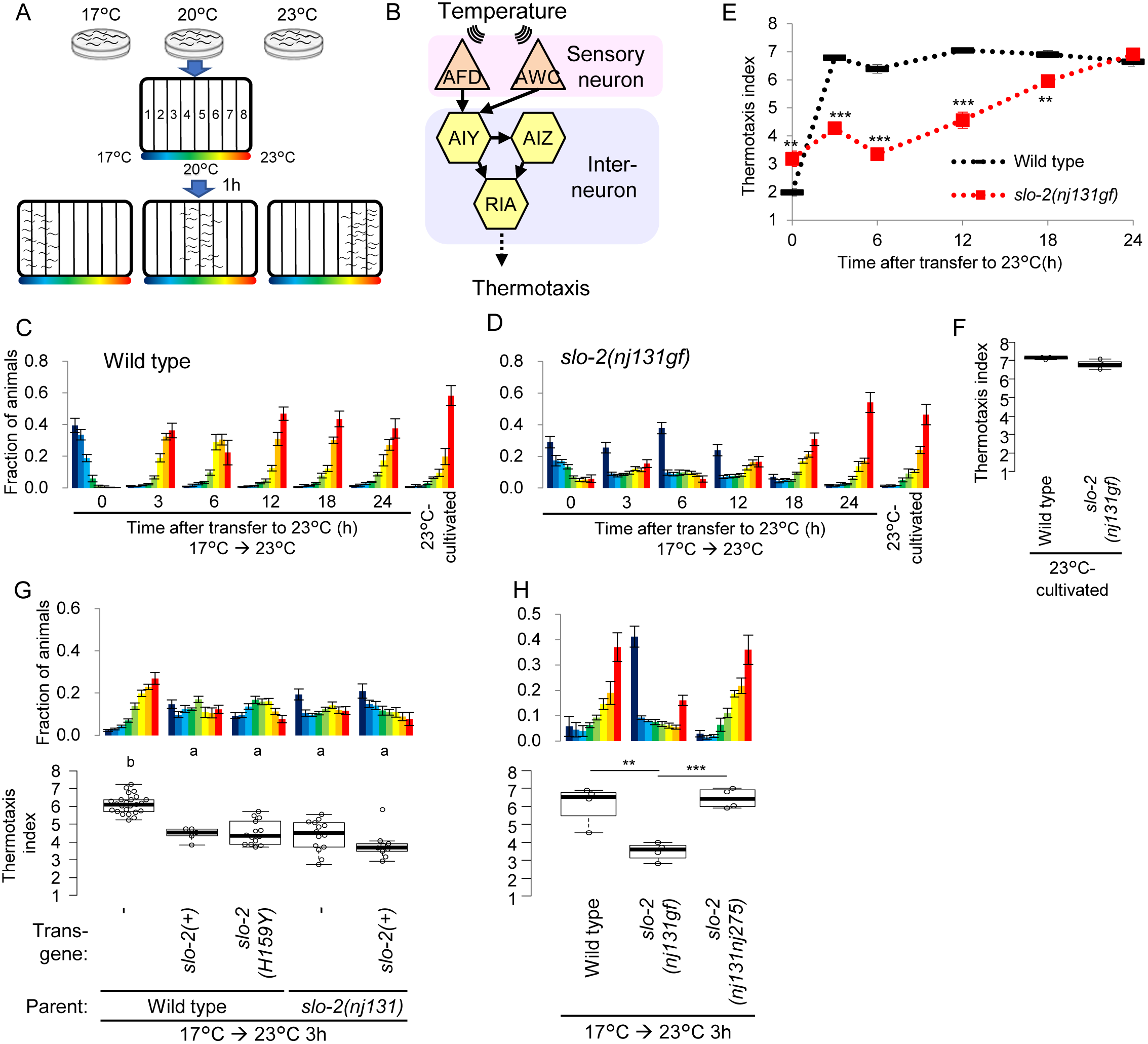
Gain of SLO-2 function decelerates transition of preference in thermotaxis behavior. (A) A scheme of thermotaxis assay. *C. elegans* cultivated at a certain temperature is placed at the center of a linear thermal gradient without food and allowed to freely migrate for 1 hour. (B) A neural circuit regulating thermotaxis. (C, D) Wild type (C) and *slo-2(nj131gf)* (D) animals were cultivated at 17°C for 5 days and then at 23°C for the time indicated or constantly at 23°C for 3 days. Animals were then placed on a thermal gradient. Numbers of animals in each section of the thermal gradient was scored, and fractions of animals were plotted in histograms. Error bars represent standard error of mean (SEM). (E) Thermotaxis indices at each time point in C and D were plotted against time after the animals were transferred to 23°C. Error bars represent SEM. **p < 0.01, ***p < 0.001 (Welch two sample t-test). (F) Thermotaxis indices for animals cultivated constantly at 23°C in C and D. (G) Genomic PCR fragments covering *slo-2* gene locus derived from either wild type or *nj131* mutant animals were injected to either wild type or *nj131* animals. Animals were cultivated at 17°C for 5 days and then at 23°C for 3 hours and then subjected to thermotaxis assay as described above. Animals carrying extra chromosomal arrays were scored for evaluating thermotaxis. Indices of strains marked with distinct alphabets differ significantly (p < 0.001) in Tukey-Kramer test. (H) Wild type, *slo-2(nj131gf)* and *slo-2(nj131nj275)* animals were cultivated at 17°C for 5 days and then at 23°C for 3 hours and then subjected to thermotaxis assay. **p < 0.01, ***p < 0.001 (Tukey-Kramer test). See also Figure S1.

The thermotaxis behavior is plastic: when cultivation temperature is shifted, animals change their temperature preference to the new cultivation temperature over the course of a few hours (Hedgecock and Russell, 1975, Figure 1C and E). AFD neurons essential for thermotaxis also acclimate to a new temperature by changing the dynamic range of responsive temperature (Biron et al., 2006; Kobayashi et al., 2016). However, the molecular mechanisms underlying the transition of the temperature preference and the AFD dynamic range are unknown.

In this study, we performed a forward genetic screen for mutants that were slow to change temperature preference in thermotaxis and demonstrated that a gain-of-function (gf) mutation in SLO-2 K^+^ channel decelerated the preference transition. The *slo-1* gene encodes the other member of the SLO family of K^+^ channel in *C. elegans*. Interestingly, the preference transition was accelerated in animals with *slo-2* and *slo-1* loss-of-function (lf) mutations. Calcium imaging of AFD revealed that SLO K^+^ channels slowed down the AFD adaptation after upshift of cultivation temperature. Moreover, through a forward genetic screen for suppressors of *slo-2gf* animals, we found that CNG-3, a subunit of CNG channels, co-operated with SLO-2 in decelerating both AFD adaptation and the transition of temperature preference.

It was recently reported that gf mutations in human SLO-2 homolog, Slack/*KCNT1*, cause early onset epilepsies (Barcia et al., 2012; Heron et al., 2012). We found that some of epilepsy-related mutations potentiated *C. elegans* SLO-2 in decelerating the transition of temperature preference. These results implicate that similar molecular and physiological mechanisms might underlie between these epilepsies and the latency regulation in *C. elegans* thermotaxis.

## Results

### Isolation of *slo-2 gf* mutant that is slow to change behavior

In order to reveal molecular mechanisms that determine the timing of behavior transition, we performed a forward genetic screen for mutants that were slow to change temperature preference after cultivation temperature was shifted (Figure S1A). One of the mutants isolated was *nj131*. When cultivated first at 17°C and then transferred to be re-cultivated at 23°C for 3 hours, wild type animals migrated to the warm region, suggesting that 3 hours of cultivation at a new temperature is sufficient for changing the temperature preference. By contrast, a majority of *nj131* animals that had been cultivated first at 17°C and then at 23°C for 3 hours still migrated to the cold region that corresponded to the old temperature (Figure 1D). Nevertheless, *nj131* mutants eventually migrate to the warm region like wild type approximately 24 hours after transferred from 17°C to 23°C (Figure 1D and E). When *nj131* mutants were cultivated constantly at 23°C, they migrated to the warm region as did wild type (Figure 1D and F), suggesting *nj131* mutants were not simply cryophilic but indeed slow to change behavior. Hereafter, we designate this abnormality of *nj131* mutants as “Slow-learning (Slw)” phenotype. When *nj131* mutants were first cultivated at 23°C and then at 17°C for 3 hours, they migrated to the cold region as did wild type (Figure S1C-E). This result suggests that *nj131* mutants are defective for preference transition specifically in response to temperature upshift.

From SNP mapping and whole-genome sequencing, *nj131* was mapped on a missense mutation in the *slo-2* gene, which alters histidine residue 159 of the gene product to tyrosine (H159Y) (Figure S1B). We then examined if *slo-2* expression can rescue the Slw phenotype in *nj131* mutants. Whereas injection of a PCR fragment containing *slo-2* locus to *nj131* animals did not rescue the Slw phenotype, introduction of mutant or wild type forms of SLO-2 in wild type genetic background phenocopied *nj131* mutants (Figure 1G), suggesting that *nj131* is a gain-of function (gf) allele of *slo-2* gene. When cultivated constantly at 23°C, animals expressing the mutant form of SLO-2 migrated to the warm region as did wild type (Figure S1F), suggesting that SLO-2 over-expression did not make animals simply cryophilic but phenocopied the *nj131* mutants. Consistent with that *nj131* is a gf allele of *slo-2, slo-2(nf101)* deletion mutants (Wei et al., 2002, Figure S1B) did not exhibit Slw phenotype and migrated to the warm region similarly to wild type when cultivated first at 17°C then at 23°C for 3 hours (Figure 3C, 3 h). Assuming that the Slw phenotype of *nj131* mutants is truly caused by gf mutation of *slo-2*, knocking out *slo-2* gene in *nj131* mutant animals should cancel the Slw phenotype. We found that knocking out *slo-2* gene by CRISPR/Cas9 clearly extinguished the Slw phenotype of *nj131* mutants (Figure 1H), indicating that the Slw phenotype of *nj131* mutants was certainly caused by gf mutation of *slo-2* gene.

**Figure 2.**
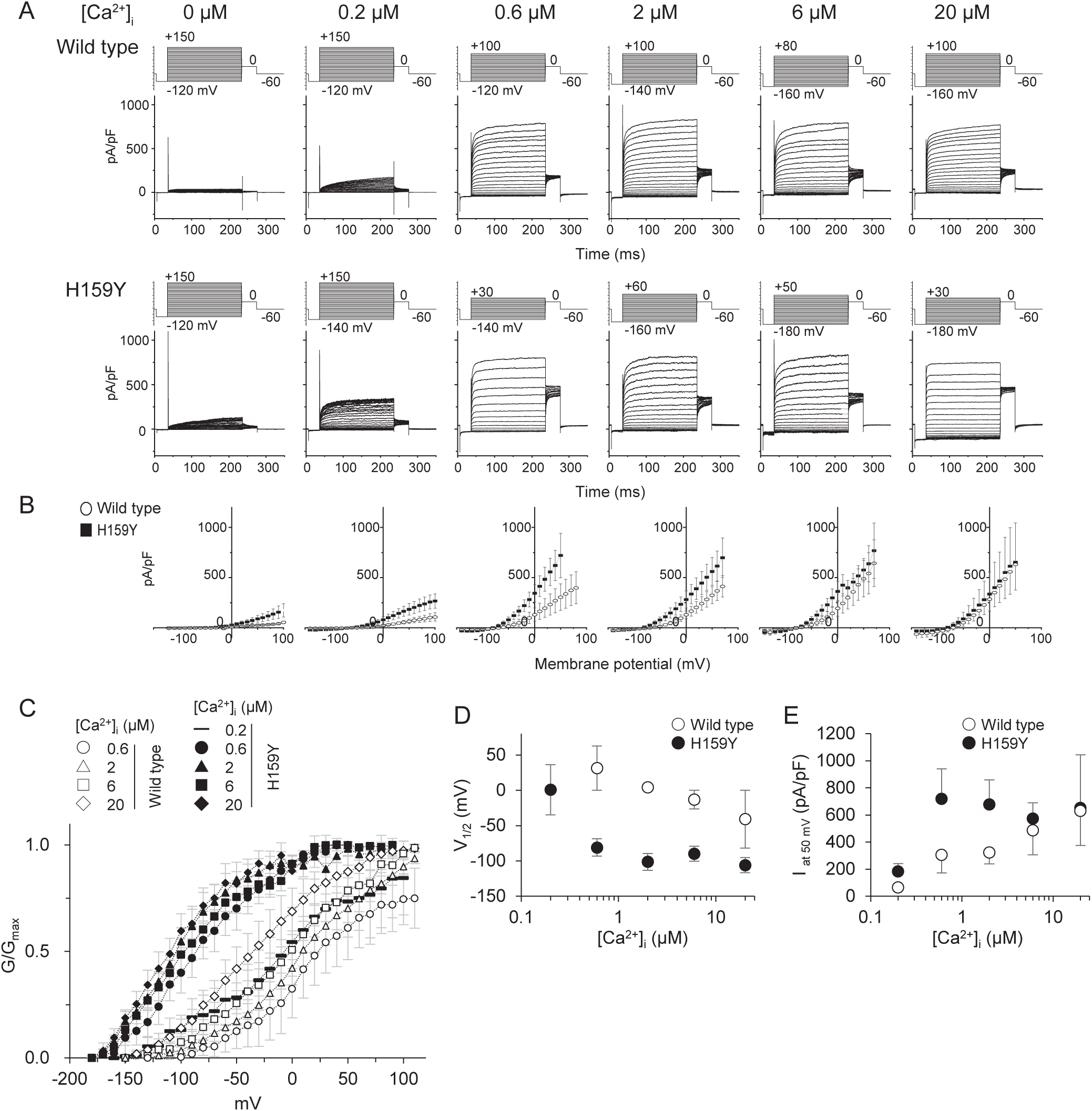
Channel currents of mutant SLO-2 (H159Y) are larger than wild type. (A) Whole cell current recordings by patch clamp method from HEK293T cells expressing either wild type or H159Y mutant SLO-2 channels. Cells were held at −60 mV and depolarizing step pulses (200 ms) were applied as indicated above the current traces. Intracellular Ca^2+^ concentration ([Ca^2+^]_i_) is indicated above the current traces. (B) Current density-voltage relationships. Current densities were measured at the end of step pulses and averaged at each membrane potential and each [Ca^2+^]_i_. (C) Conductance (G/G_max_)-voltage (V) relationships. Conductance (G) was measured as the tail current amplitudes at 0 mV which follows various step pulse stimulations, and was normalized to the recorded maximal one at most depolarized potential. The meaning of symbols is as indicated in the figure. (D) Voltage at half-maximal activation (Vi/_2_) of wild type (open circles) and the H159Y mutant (filled circles) channels calculated from K was plotted against [Ca^2+^]_i_. (E) Current density values at 50mV plotted as a function of [Ca^2+^]_i_. Current density at the depolarization to 50 mV pulse was measured and averaged in [Ca^2+^]_i_ and then plotted against [Ca^2+^]_i_.

**Figure 3.**
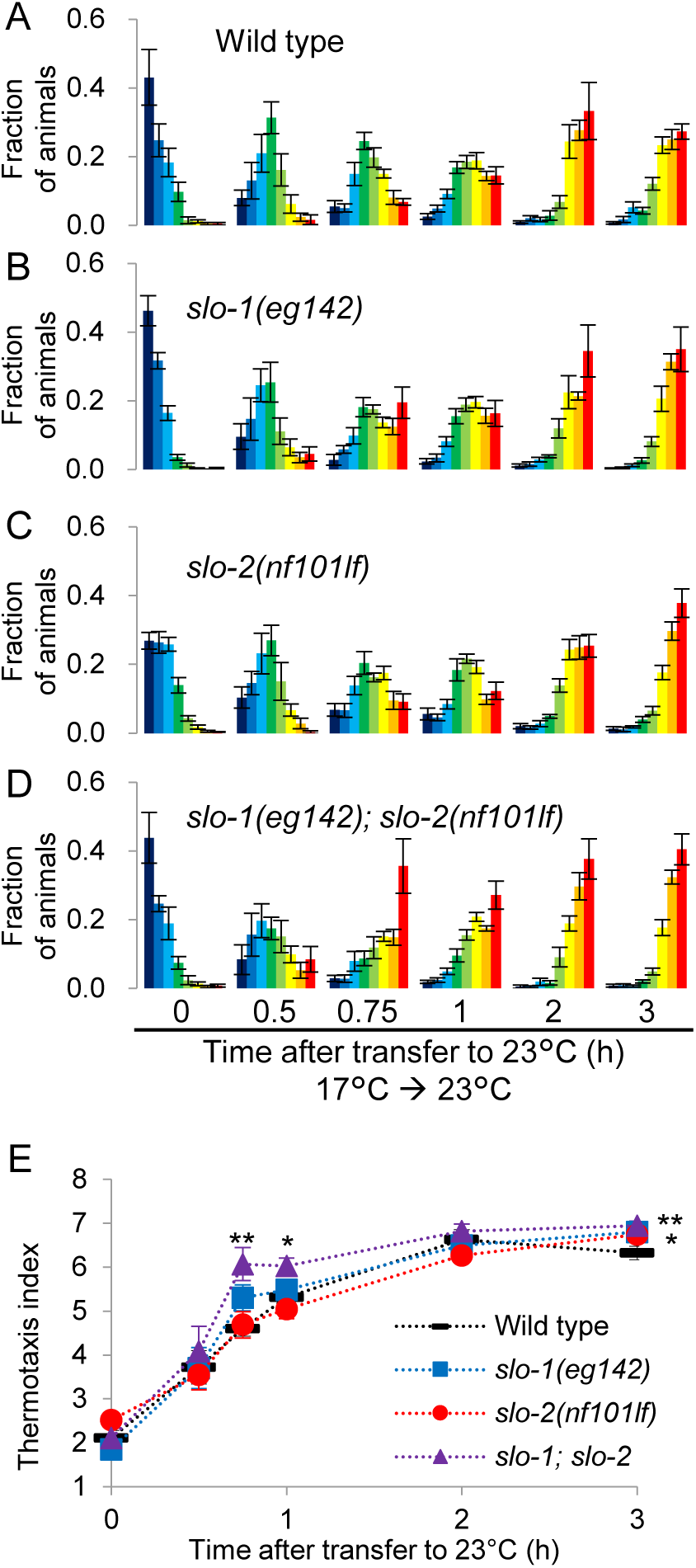
*slo-1; slo-2* double loss-of -function mutants are fast to change behavior. (A-D) Wild type (A), *slo-1(eg142)* (B), *slo-2(nf101)* (C) and *slo-1(eg142); slo-2(nf101)* (D) animals were cultivated at 17°C for 5 days and then at 23°C for the time indicated. Animals were then subjected to thermotaxis assay. Fractions of animals were plotted in histograms. (E) Thermotaxis indices were plotted against time after animals were transferred to 23°C. *p < 0.05, **p < 0.01 (Dunnett test against wild type). See also Figure S1, S2 and S3.

*slo-2* gene encodes a Ca^2+^-dependent K^+^ channel, which consists of an N-terminal transmembrane domain and a large C-terminal intracellular regulatory domain (Yuan et al., 2000). The H159Y mutation in *nj131* mutants is within the third helix (S3) of transmembrane domain (Hite et al., 2015, Figure S2). To examine the effects of the H159Y mutation on the function of SLO-2 channel, we recorded whole cell currents by patch clamp method from HEK293T cells expressing either wild type or H159Y mutant of SLO-2 channels. As has been reported (Yuan et al., 2000), depolarization failed to evoke increases in outward currents of wild type and mutant channels when [Ca^2+^]_i_ was zero, but elicited them when [Ca^2+^]_i_ was more than 0.6 μM (Figure 2A and B). With [Ca^2+^]_i_Of 0.2 μM, the H159Y mutant was clearly activated in a membrane potential, -dependent manner but wild type was activated only slightly. The conductance-membrane potential (GV) relationship was shifted in both channels towards hyperpolarized potentials with the increase in [Ca^2+^]_i_. In the same [Ca^2+^]_i_, the GV relationship of the H159Y mutant was located at more hyperpolarized potential than wild type, showing a higher sensitivity to [Ca^2+^]_i_ and to depolarization (Figure 2C-E). These results demonstrate that the activity of the SLO-2 channel increased by the H159Y point mutation and proved that this mutation causes gain of function in *slo-2*.

### Animals lacking both *slo-1* and *slo-2* are fast to change behavior

Since a gf mutation in *slo-2* decelerated the transition of temperature preference, we next examined if the loss of *slo-2* accelerated it. Since the preference transition was not accelerated in *slo-2(nf101)* deletion mutants (Figure 3C and E), we examined animals lacking both *slo-2* and *slo-1*. SLO-1 is structurally homologous to SLO-2 (Figure S2 and S3A), and SLO-1 and SLO-2 channels constitute the SLO family of K^+^ channel in *C. elegans*. In *slo-1(eg142); slo-2(nf101)* double lf mutants, preference transition was slightly but significantly accelerated 45 min and 1 h after upshift of cultivation temperature (Figure 3D and E). Another double lf mutant *slo-1(gk262550); slo-2(ok2214)* also showed accelerated preference transition (Figure S3C and D). These results suggest that endogenous SLO K^+^ channels function redundantly (Alqadah et al., 2016) in response to upshift of cultivation temperature. After temperature downshift, *slo-1(eg142); slo-2(nf101)* double lf mutants changed the preference no later than wild type (Figure S3E), suggesting again that SLO K^+^ channels regulate the preference transition specifically when temperature is upshifted. Although vertebrate homologs of SLO-2 and SLO-1 were shown to interact (Joiner et al., 1998), *slo-2(nj131gf)* mutation decelerated the preference transition even in the absence of SLO-1 (Figure S3F), indicating that the gf mutant form of SLO-2 can act independently of SLO-1.

### SLO K^+^ channels act in AFD thermosensory neuron

Since *slo-2* is expressed broadly in neurons and muscles (Figure S4A, Yuan et al., 2000), we aimed to find the site(s) of action of SLO K^+^ channels during the transition of temperature preference. Expressing SLO-2 pan-neuronally or specifically in AFD thermosensory neuron phenocopied *slo-2(nj131gf)* mutants, while expression in AWC chemo- and thermosensory neuron, in interneurons involved in regulation of thermotaxis or in body wall muscles did not (Figure 4A). These results suggest that SLO-2 acts in AFD to decelerate the preference transition.

**Figure 4.**
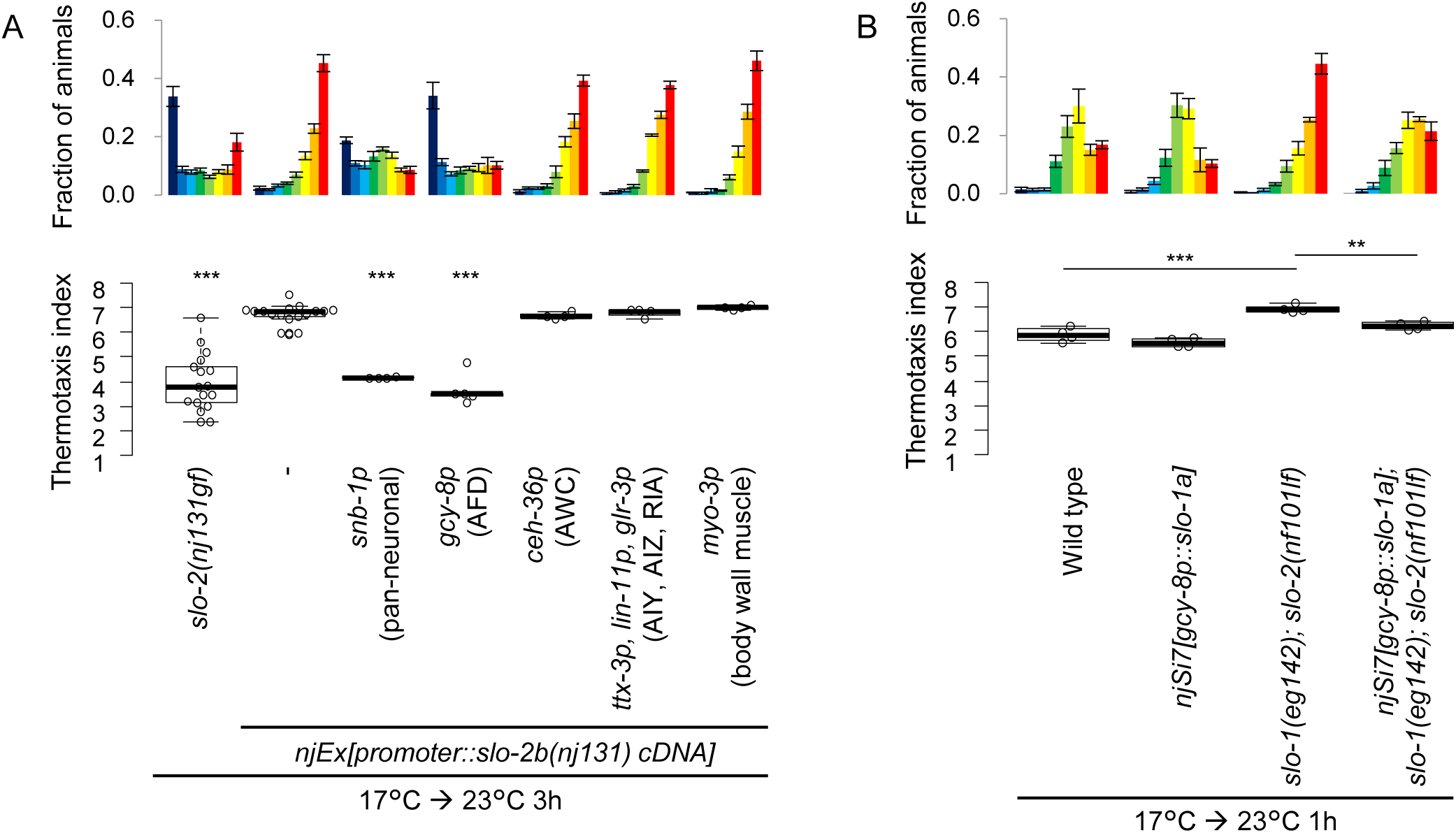
SLO K* channels act in AFD thermosensory neuron to decelerate the preference transition in thermotaxis. (A) Animals expressing H159Y mutant form of SLO-2 isoform b under the control of promoters indicated were cultivated at 17°C for 5 days and then at 23°C for 3 hours and subjected to thermotaxis assay. Fractions of animals and thermotaxis indices were plotted. ***p < 0.001 (Dunnett test against wild type). (B) Wild type animals, wild type animals that express SLO-1 in AFD from a single copy of transgene inserted in a genome, *slo-1(eg142); slo-2(nf101)* animals and *slo-1(eg142); slo-2(nf101)* animals that express SLO-1 in AFD from a single copy insertion were cultivated at 17°C for 5 days and then at 23°C for 1 hour. Animals were then subjected to thermotaxis assay. **p < 0.01, ***p < 0.001 (Tukey-Kramer test). See also Figure S4.

Abilities of wild type and the H159Y mutant form of SLO-2 to decelerate the preference transition were compared by expressing each form in AFD with various concentrations. Injection of plasmid encoding the H159Y form at 1 ng/μl decelerated the preference transition more than injection of wild type at 20 ng/μl, suggesting that H159Y form is far more potent to decelerate the preference transition (Figure S4B and C). These results are consistent with the higher activity of mutant channel revealed by electrophysiological experiments (Figure 2C-E).

We next examined if the accelerated preference transition in *slo-1; slo-2* double lf mutants were caused by the loss of SLO channels in AFD. Expressing SLO-1 in AFD rescued the accelerated preference transition in *slo-1; slo-2* mutants (Figure 4B), indicating that loss of SLO-1 is responsible for the accelerated preference transition and that SLO-1 acts in AFD. Taken together with the results in Figure 4A, SLO K^+^ channels were revealed to act in AFD to decelerate the preference transition.

### CNG-3 cyclic nucleotide-gated channel functions together with SLO-2

To investigate how SLO K^+^ channels decelerate the preference transition, we performed a genetic screen for suppressors of *slo-2(nj131gf)* mutants in order to isolate molecules that functionally interact with SLO-2 (Figure S5A). Of mutagenized *slo-2(nj131gf)* mutant animals, *nj172* that partially suppressed the Slw phenotype of *slo-2(nj131gf)* was isolated (Figure 5D and G). *nj172; slo-2(nj131gf)* double mutants migrated to the cold region as did wild type when cultivated constantly at 17°C (Figure 5D, 0 h), suggesting that *nj172* did not make animals simply thermophilic but suppressed the Slw phenotype of *slo-2(nj131gf)* animals.

**Figure 5.**
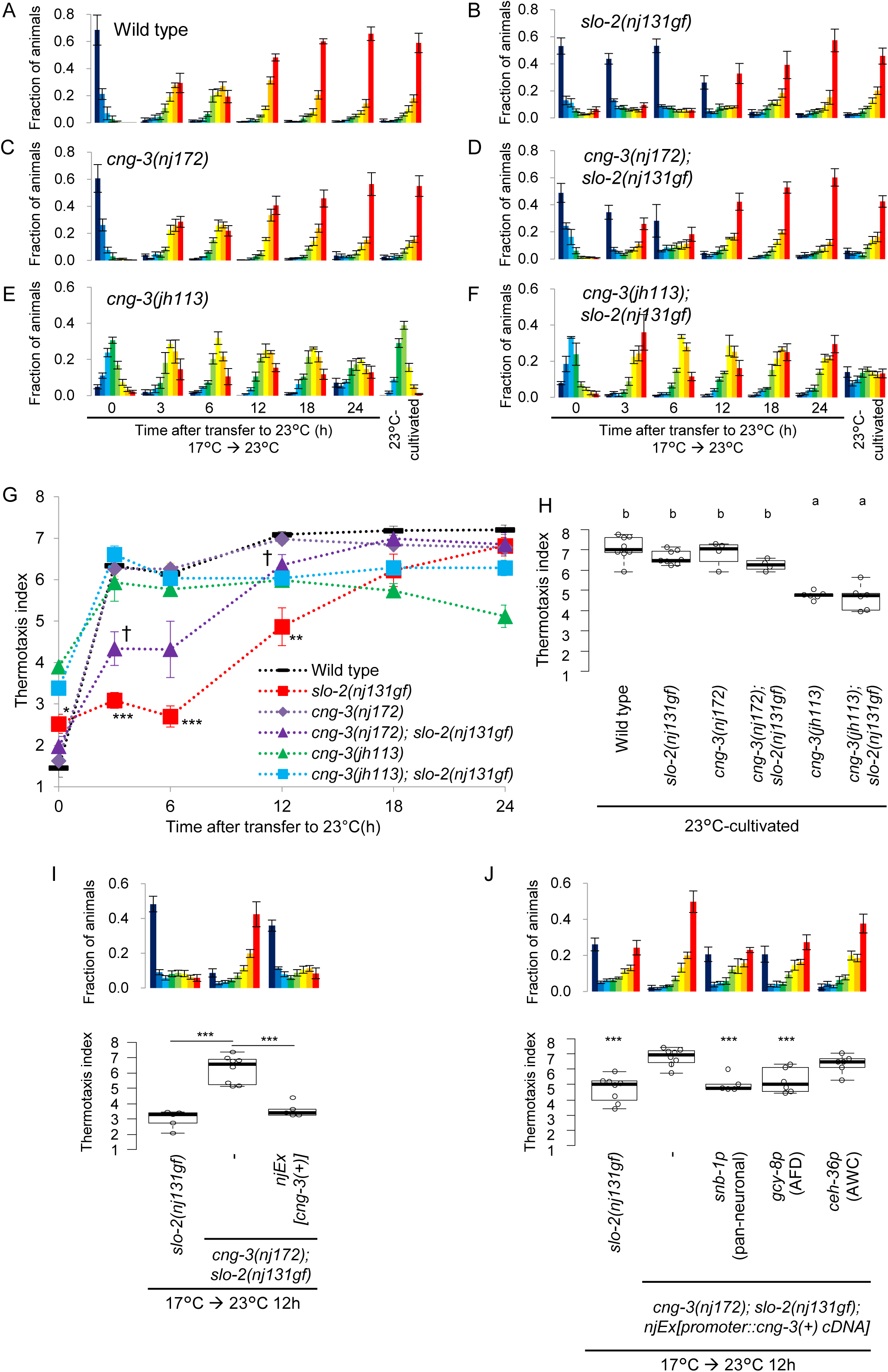
*cng-3* loss-of-function suppresses the decelerated preference transition in *slo-2(nj131gf)* mutants. (A-F) Wild type (A), *slo-2(nj131gf)* (B), *cng-3(nj172)* (C), *cng-3(nj172); slo-2(nj131gf)* (D), *cng-3(jh113)* (E) and *cng-3(jh113); slo-2(nj131gf)* (F) animals were cultivated at 17°C for 5 days and then at 23°C for the time indicated or constantly at 23°C for 3 days. Animals were then subjected to thermotaxis assay. Fractions of animals and thermotaxis indices were plotted. (G) Thermotaxis indices at each time point in A-F were plotted against time after the animals were transferred to 23°C. Error bars represent SEM. *p < 0.05, **p < 0.01, ***p < 0.001 between wild type and *slo-2(nj131gf)*, and +p < 0.05 between *slo-2(nj131gf)* and *cng-3(nj172); slo-2(nj131gf)* (Tukey-Kramer test). (H) Thermotaxis indices for animals cultivated constantly at 23°C in A-F. Indices of strains marked with distinct alphabets differ significantly (p < 0.001) in Tukey-Kramer test. (I) A genomic PCR fragments covering the *cng-3* locus was injected to *cng-3(nj172); slo-2(nj131gf)* animals. Animals were cultivated first at 17°C then at 23°C for the time indicated and subjected to thermotaxis assay. *p < 0.05, **p < 0.01, ***p < 0.001 (Tukey-Kramer test). (J) *cng-3(nj172); slo-2(nj131gf)* animals expressing *cng-3* under the control of promoters indicated were cultivated at 17°C for 5 days and then at 23°C for 12 hours and subjected to thermotaxis assay. ***p < 0.001 (Dunnett test against *cng-3(nj172); slo-2(nj131gf)* animals). See also Figure S5.

SNP mapping and subsequent rescue experiments showed that *nj172* is an allele of *cng-3* (Figure 5I, S5H and I). The *cng-3* gene encodes a subunit of cyclic nucleotide-gated (CNG) cation channel. *nj172* mutants carried a missense mutation in the *cng-3* gene, which altered methionine residue 467 to isoleucine (M467I) within a conserved C-terminal cyclic nucleotide-binding domain of the gene product (Kaupp and Seifert, 2002). When cultivated constantly at 23°C, *nj172; slo-2(nj131gf)* animals expressing *cng-3* migrated to the warm region as did wild type (Figure S5I), suggesting that *cng-3* expression did not make animals simply cryophilic but canceled the suppression by *nj172*.

We also characterized *cng-3(jh113)* deletion mutants. When first cultivated at 17°C and then at 23°C for 3 hours, *cng-3(jh113); slo-2(nj131gf)* animals migrated toward the warm region on a thermal gradient (Figure 5F and G), suggesting that *cng-3(jh113)* deletion mutation also suppressed the Slw phenotype of *slo-2(nj131gf)* animals. The extent of suppression by *cng-3(jh113)* deletion mutation seemed stronger than that by *cng-3(nj172)* mutation, suggesting that *cng-3(nj172)* is a reduction-of-function allele. When cultivated constantly at 17°C or at 23°C, both *cng-3(jh113); slo-2(nj131gf)* and *cng-3(jh113)* mutants did not show clear preference for the cultivation temperature (Figure 5E-H, Smith et al., 2013). These results indicated that CNG-3 is necessary for thermotaxis after cultivation under constant temperature but dispensable for acquisition of a novel preference after cultivation temperature is shifted. Other CNG subunits TAX-2 and TAX-4, however, are critial for thermosensation, since animals lacking either TAX-2 or TAX-4 were completely athermotactic regardless of whether cultivation temperature was shifted or not (Figure S5J).

CNG-3 was reported to be expressed in sensory neurons including AFD (Cho et al., 2004; Wojtyniak et al., 2013). To determine the site(s) where CNG-3 acts, we expressed CNG-3 in *cng-3(nj172); slo-2(nj131gf)* mutants cell-specifically. Expression of CNG-3 in AFD but not in AWC canceled the suppression by *cng-3(nj172)* (Figure 5J), suggesting that CNG-3 functions in AFD as do SLO channels (Figure 4).

### SLO and CNG channels slowed down the adaptation of AFD

Since SLO-2 was shown to act in AFD thermosensory neuron to decelerate the transition of temperature preference, we next examined whether SLO-2 acts upstream of Ca^2+^ response in AFD by Ca^2+^imaging. AFD responds to temperature rise above the past cultivation temperature (Kimura et al., 2004; Kobayashi et al., 2016). In wild type animals cultivated constantly at 17°C, AFD started increasing Ca^2+^ level approximately at 15°C in response to a temperature ramp from 14°C to 24°C (Figure 6A and G). The onset temperature of AFD responses in wild type animals cultivated at 23°C was around 23°C, and that in animals cultivated first at 17°C then at 23°C for 3 hours was around 21°C. Although we did not detect any obvious abnormality in the onset temperature of *slo-2(nj131gf)* animals cultivated constantly at 17°C or at 23°C, the onset temperature in *slo-2(nj131gf)* animals cultivated first at 17°C and then at 23°C for 3 hours was significantly lower than that in wild type (Figure 6B and G). When cultivated longer at 23°C, the onset temperature in *slo-2(nj131gf)* animals gradually changed to the level comparable to that in wild type (Figure 6H). These results suggest that *slo-2(nj131gf)* mutation slowed down transition of AFD dynamic range, i.e. adaptation of AFD. This adaptation of AFD preceded the behavioral transition both in wild type and in *slo-2(nj131gf)* mutants (Figure 1E and 6H). These results implicate a mechanism acting either downstream of Ca^2+^influx in AFD or in downstream neural circuits, which reflects the transition of AFD dynamic range on thermotaxis behavior with delay. Moreover, expression of SLO-2 specifically in AFD neurons also lowered the onset temperature in animals cultivated first at 17°C and then at 23°C for 3 hours (Figure 6E and G), suggesting that enhanced SLO-2 channel currents in AFD slow down the AFD adaptation cell-autonomously.

**Figure 6.**
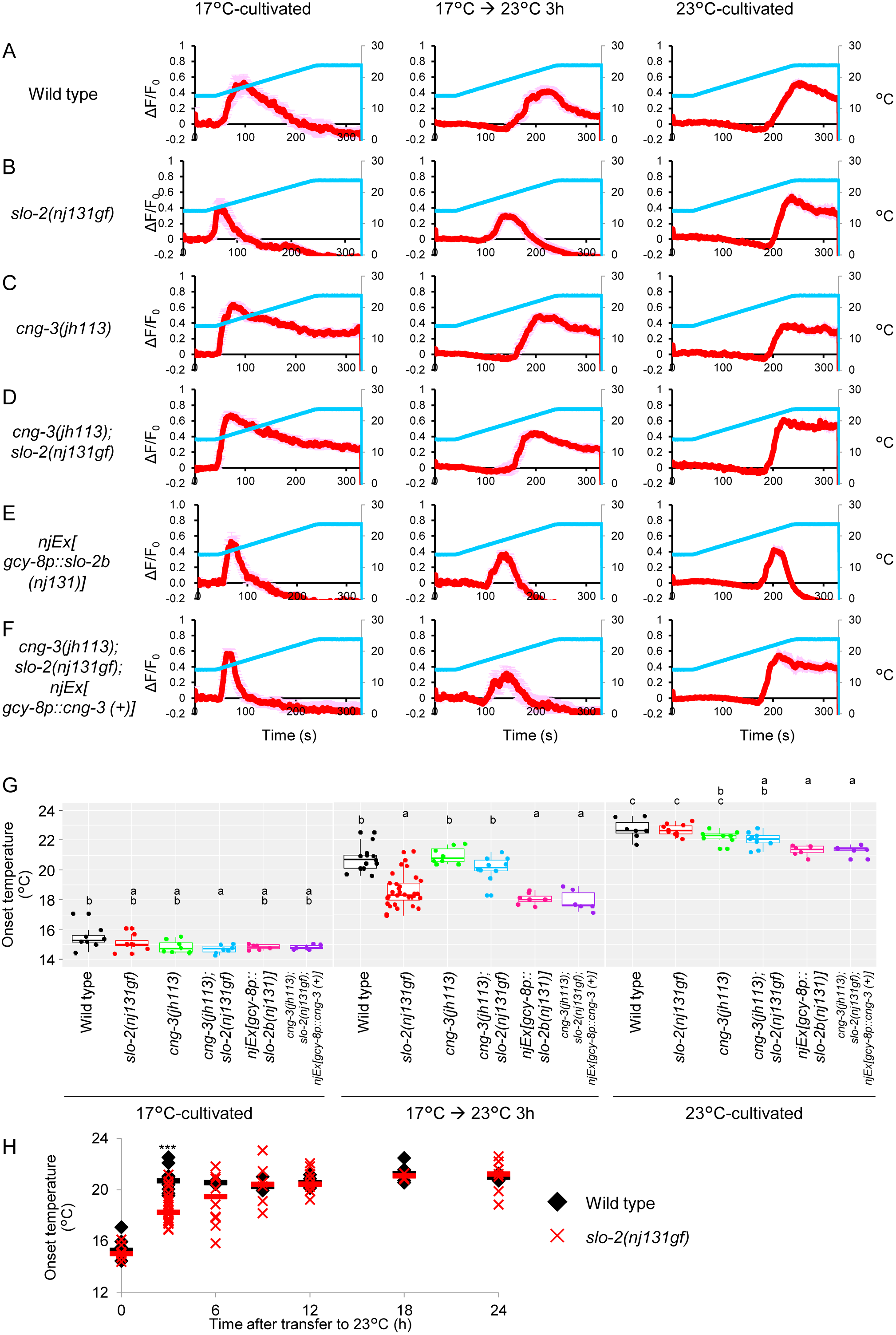
SLO K* channels slow down AFD adaptation. (A-F) Wild type (A), *slo-2(nj131gf)* (B), *cng-3(jh113)* (C) and *cng-3(jh113); slo-2(nj131gf)* (D) animals, animals expressing SLO-2b(H159Y) in AFD (E) and *cng-3(jh113); slo-2(nj131gf)* animals expressing CNG-3 in AFD (F) that express R-CaMP2 in AFD were cultivated either at 17°C for 5 days, at 17°C for 5 days then at 23°C for 3 hours, or at 23°C for 3 days. Animals were then subjected to Ca^2+^ imaging analysis. The animals were initially kept at 14°C for 40 seconds and then subjected to a linear temperature rise to 24°C over 200 seconds followed by 90 seconds of incubation at 24°C. Temperature presented and changes in fluorescence intensity were indicated with blue and red lines, respectively. (G) Onset temperature, at which the response of AFD reached the half of the maximum, during the experiment above (A-F) was plotted. Onset temperature of strains marked with distinct alphabets differ significantly (p < 0.05) in Tukey-Kramer test within the same cultivation condition. (H) Wild type and *slo-2(nj131gf)* animals that express R-CaMP2 in AFD were cultivated at 17°C for 5 days then at 23°C for the time indicated. Animals were then subjected to Ca^2+^ imaging with the same ramp temperature stimulus as in A-F. Onset temperature was plotted, and medians were indicated with bars. ***p < 0.001 (Welch two sample t-test).

Since the Slw phenotype in *slo-2(nj131gf)* mutants was suppressed by *cng-3* mutation, we next examined whether the slowed AFD adaptation after upshift of cultivation temperature in *slo-2(nj131gf)* mutants was also suppressed by *cng-3* mutation. When cultivated first at 17°C and then at 23°C for 3 hours, *cng-3(jh113)* suppressed the lowered onset temperature of AFD in *slo-2(nj131gf)* animals to the level comparable to that in wild type (Figure 6D and G). Moreover, AFD-specific expression of CNG-3 in *cng-3(jh113); slo-2(nj131gf)* double mutants canceled the suppression of decelerated adaptation by *cng-3* (Figure 6F and G). These results suggest that CNG-3 functionally interacts with SLO-2 in AFD to slow down the AFD adaptation and thereby to decelerate behavioral transition. Although *cng-3(jh113)* mutants showed aberrant thermotaxis behavior when cultivated constantly at 17°C or at 23°C (Figure 5E), AFD in *cng-3(jh113)* mutants responded to temperature rise at around cultivation temperature similarly to wild type, except that decrease of Ca^2+^level was slower (Figure 6C). This is in a sharp contrast with *tax-2* or *tax-4* mutants where AFD is totally silent (Kobayashi et al., 2016; Ramot et al., 2008).

We next examined how endogenous SLO channels affect the timing of changes in AFD dynamic range. AFD changes its dynamic range within a few hours after cultivation temperature is shifted (Biron et al., 2006; Yu et al., 2014), whereas AFD does not respond to temperature rise beyond the temperature that is far high as compared to the cultivation temperature (Kobayashi et al., 2016). Thus, the animals cultivated at 17°C were investigated when their AFD regain responses after temperature upshift, using a temperature program that contained short-term temperature oscillations with a period of 20 secs (0.05 Hz) around 17°C followed by a shift to 23°C, which mimicked the upshift of cultivation temperature, and long-term oscillations around 23°C (Figure 7A-E). AFD in wild type animals showed phase-locked responses to the oscillations around 17°C, a large response to the temperature upshift and a gradual Ca^2+^ decrease and then faintly regained phase-locked responses to the oscillations around 23°C in 20 minutes after the upshift (Figure 7A). In contrast, AFD in *slo-2(nj131gf)* animals showed a sharp Ca^2+^ decrease and a profound silencing after the temperature upshift and did not regain the phase-locked response in 20 minutes (Figure 7B). On the other hand, AFD in *slo-1; slo-2* double if mutants showed clear phase-locked responses to the oscillations from earlier time point than wild type (Figure 7E). Fourier transform of the AFD responses further supported these observations (Figure 7F-J). Fourier transform of the temperature record exhibited a clear peak at 0.05 Hz. The peak at 0.05 Hz in AFD responses of *slo-1; slo-2* double if mutants was much more explicit than that of wild type animals (Figure 7F and J). In time courses of Fourier transform, component around 0.05 Hz appeared earlier in *slo-1; slo-2* double if mutants than in wild type (Figure 7K and O). Taken together, these results further support that SLO K^+^ channels slow down the AFD adaptation after upshift of cultivation temperature.

**Figure 7.**
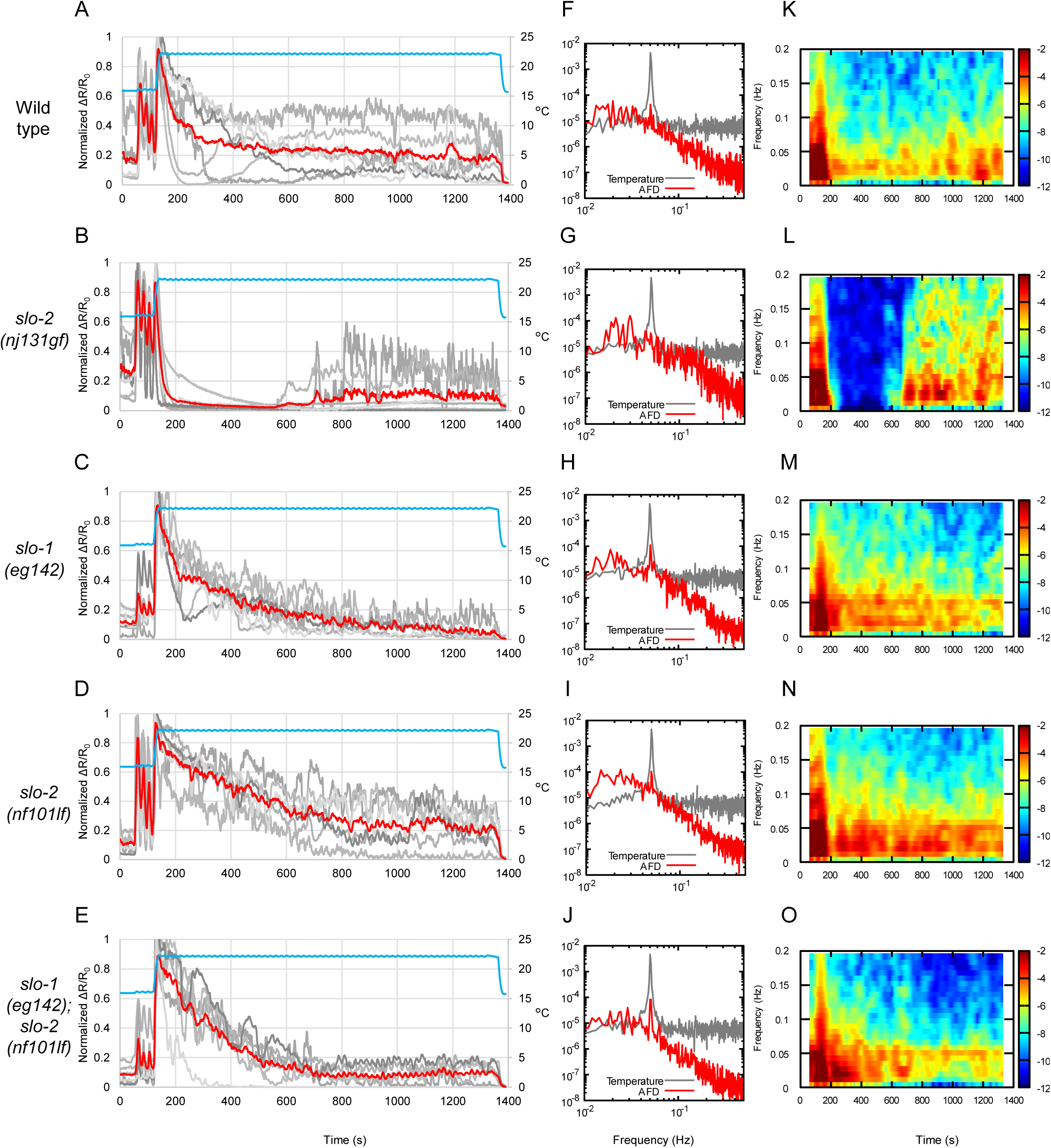
Endogenous SLO K* channels slow down AFD adaptation. (A-F) Wild type (A), *slo-2(nj131gf)* (B), *slo-1(eg142)* (C), *slo-2(nf101)* (D) and *slo-1(eg142); slo-2(nf101)* (E) animals that express GCaMP3 and tagRFP in AFD were cultivated at 17°C for 5 days and subjected to Ca^2+^ imaging analysis with a temperature stimulus containing oscillations with a period of 20 secs (0.05 Hz) around 17°C followed by temperature upshift to 23°C and oscillations around 23°C. Intensity of green fluorescence was divided by that of red fluorescence, and the ratio was normalized and plotted against time. Gray and red lines indicate traces from individual animals and average, respectively. (F-J) Fourier transform was computed with the Hanning window on the temperature program and the ratio of fluorescence intensity between 401 and 1390 sec of each animal in A-E. Averaged power spectra were plotted against frequency. (K-O) From the ratio of fluorescence intensity of each animal in A-F, trends detected by Butterworth filter were removed. The resulting signals were divided into segments of 128 sec, and the Fourier transform was separately computed on each segment. Time evolutions of averaged spectrograms were plotted in a color map against centric time of each segment.

### Epilepsy-related mutations potentiate SLO-2 in decelerating the transition of temperature preference

Gain-of-function mutations in *slo-2* human homolog, *Slack/KCNT1* that encodes a Na^+^-gated K^+^ channel mainly expressed in brains (Yuan et al., 2003), cause early onset epilepsies such as malignant migrating partial seizures of infancy (MMPSI) and autosomal dominant nocturnal frontal lobe epilepsy (ADNFLE) (Barcia et al., 2012; Heron et al., 2012). Epilepsy-related mutants Slack channel have enhanced channel activity (Barcia et al., 2012; Kim et al., 2014a; Milligan et al., 2014; Tang et al., 2016). Analogously, H159Y mutant form of *C. elegans* SLO-2 isolated in this study had higher channel activity (Figure 2), suggesting that both human epilepsies and the decelerated transition of temperature preference in thermotaxis are caused by excess K^+^efflux through SLO-2 channels. Therefore, we tested whether introduction of the epilepsy-related mutations in *C. elegans* SLO-2 (Figure S2) could phenocopy the gf nature of H159Y in decelerating the preference transition in thermotaxis.

We introduced each of six epilepsy-related mutations into *C. elegans slo-2* DNA and injected each DNA into *slo-2(nf101)* deletion mutants. Animals expressing wild type or mutant forms of SLO-2 in AFD were cultivated first at 17°C then at 23°C for 3 hours and were subjected to thermotaxis assay. Expression of two out of six epilepsy-related mutant SLO-2 decelerated the preference transition more than that of wild type did (Figure 8A and B). These results implicate that common molecular mechanisms might underlie between human epilepsies and the preference transition in *C. elegans* thermotaxis and that at least these two mutations potentiate SLO-2/Slack channel regardless of their differences such as gating either by Ca^2+^ or Na^+^. We also found that the *cng-3* mutation suppressed the decelerated preference transition caused by expressing SLO-2(R376Q) in AFD (Figure 8C), implying that the inhibition of CNG channels might attenuate epilepsy symptoms. It is of clinical interest to examine whether *Slack(gf)* rodent models show epilepsy-related phenotypes, and inhibitors for CNG inhibitors attenuate those phenotypes.

**Figure 8.**
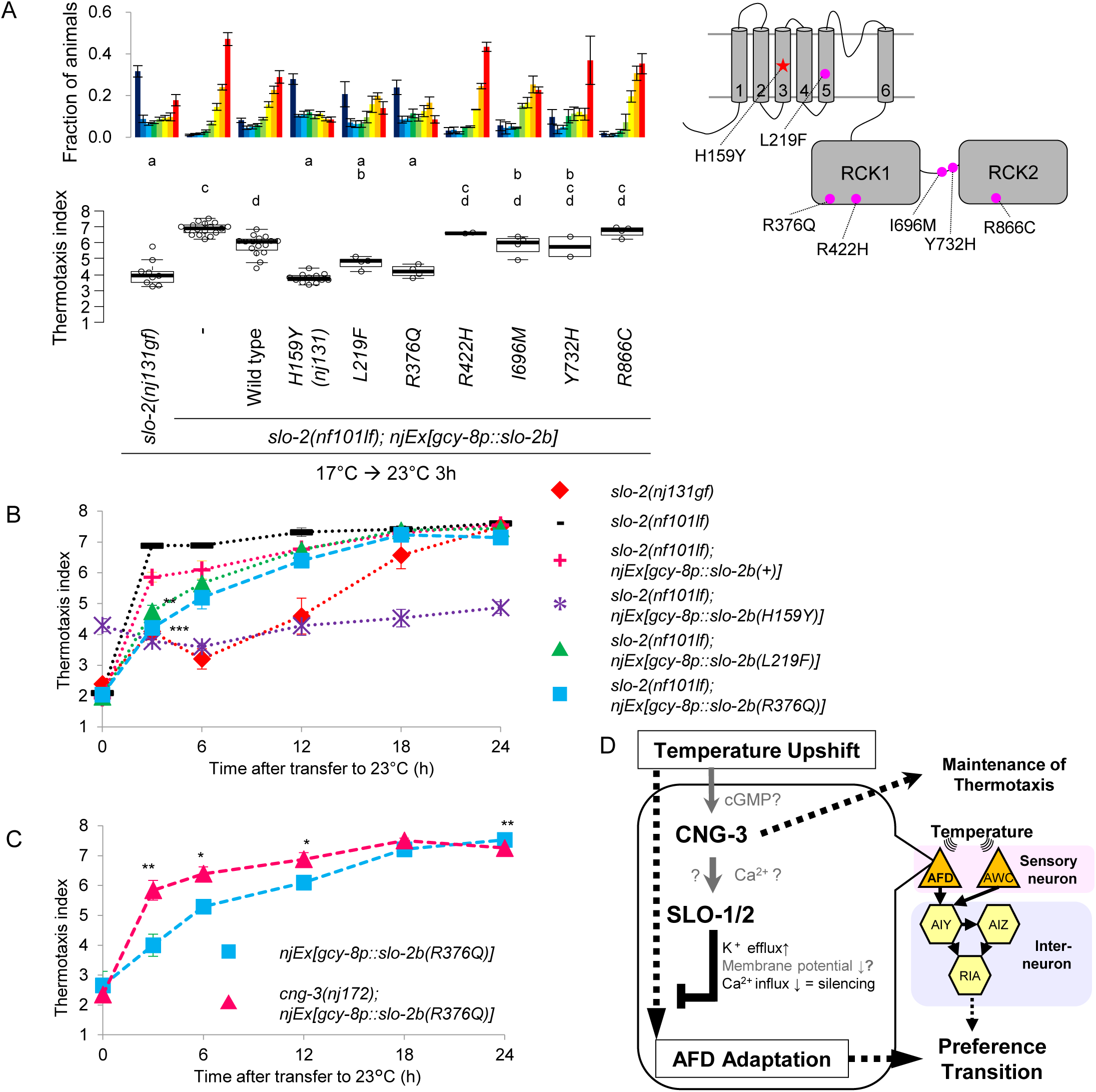
Epilepsy-related mutations potentiate SLO-2. (A) *slo-2(nf101)* animals expressing either wild type or indicated mutant form of SLO-2b in AFD were cultivated at 17°C for 5 days and then at 23°C for 3 hours and then subjected to thermotaxis assay. Indices of strains marked with distinct alphabets differ significantly (p < 0.05) in Tukey-Kramer test. (B) *slo-2(nf101)* animals expressing either wild type or indicated mutant form of SLO-2b in AFD were cultivated at 17°C for 5 days and then at 23°C for the time indicated. Animals were then subjected to thermotaxis assay. **p < 0.01, ***p < 0.001 (Tukey-Kramer test, compared with animals expressing SLO-2b(+)). See also Figure S2, S3 and S6. (C) Animals expressing SLO-2b(R376Q) in AFD on either wild type or *cng-3(nj172)* background were cultivated at 17°C for 5 days and then at 23°C for the time indicated. Animals were then subjected to thermotaxis assay. *p < 0.05, **p < 0.01 (Welch two sample t-test). (D) A model for regulation of the timing of the preference transition in thermotaxis. SLO K^+^ channels and CNG-3 generate latency for the preference transition in thermotaxis after upshift of cultivation temperature by acting in AFD to slow down the AFD adaptation to new temperature. See also Figure S2, S3, S6 and S7.

## Discussion

In this study, we demonstrated that SLO K^+^ channels function together with CNG-3 channel in AFD thermosensory neuron to slow down the transition of AFD dynamic range and thereby generate latency for transition of the temperature preference in thermotaxis upon temperature upshift. In response to upshift of ambient temperature and resulting large Ca^2+^ influx into AFD, K^+^ efflux through SLO channels would repolarize membrane potential and decrease [Ca^2+^]_i_, which would antagonize premature AFD adaptation (Figure 8D). Although expression levels of AFD-specific guanylyl cyclases are shown to increase when cultivation temperature is upshifted (Yu et al., 2014), this induction was unaffected by *slo-2(nj131gf)* mutation (Figure S7A-C). We also found a molecular and physiological link between early onset epilepsies and the preference transition in thermotaxis, providing a model system useful for understanding pathogenesis of these epilepsies and for drug screening.

Although CNG channels are generally thought to depolarize membrane potential (Kaupp and Seifert, 2002), our results implied that CNG-3 acts with SLO K^+^ channels to hyperpolarize AFD (Figure 5). Considering that SLO-1 and SLO-2 of *C. elegans* are both gated by Ca^2+^ (Salkoff et al., 2006) and that CNG channels are generally permeable to Ca^2+^ (Kaupp and Seifert, 2002), it would be reasonable to hypothesize that SLO and CNG channels are co-localized in a limited region, where local Ca^2+^ influx through CNG channels including CNG-3 activates SLO K^+^ channels.

While the onset temperature of AFD in *slo-2(nj131gf)* animals cultivated at 17°C was comparable with that in wild type, Ca^2+^ level decreased sharply and to beneath the base line in *slo-2(nj131gf)* animals (Figure 6B). This result is consistent with the important roles of SLO-2 in post-excitatory repolarization of membrane potential in body wall muscles (Santi et al., 2003; Yuan et al., 2003; Liu et al., 2011) and in motor neurons (Liu et al., 2014) in *C. elegans* as well as in mammalian neurons (Franceschetti et al., 2003; Gao et al., 2008). Given that AFD-specific SLO-2 expression also altered the AFD response and thermotaxis behavior of animals cultivated at 17°C in a manner similar to *slo-2(nj131gf)* mutants (Figure 6E and S6D, 0 h and 6G), the sharp Ca^2+^ decrease in AFD response might result in a slightly diffused distribution of the *slo-2(nj131gf)* animals cultivated at 17°C on a thermal gradient (Figure 1D). The sharp Ca^2+^ decrease in AFD response and the diffused distribution of *slo-2(nj131gf)* animals on a thermal gradient were also suppressed by *cng-3* mutation (Figure 5D and 6D), further supporting their causal relationship.

Although a number of studies so far have focused on alteration of relationships between unconditional stimuli (US) and conditional stimuli (CS) (reviewed in Aoki et al., 2017), our study focused on CS transition. While *C. elegans* animals associate CS such as temperature or chemicals with preferable US such as the existence of food and are attracted to that CS, the animals also associate CS with unpreferable US and are no longer attracted to or even avoid that CS. For instance, the role of insulin-PI3K-Akt pathway has been intensively studied during association between CS with starvation as a negative US (Chalasani et al., 2010; Kodama et al., 2006; Tomioka et al., 2006). Meanwhile, mechanisms underlying decay of the US-CS association have also been described (Inoue et al., 2013; Hadziselimovic et al., 2014; Vukojevic et al., 2012). In this study, we performed a forward genetic screen that focused on CS transition, and newly identified molecules whose roles in learning had been previously undescribed. Our results suggested that SLO and CNG channels act together to modulate the transition of dynamic range of a sensory neuron, which can encode the CS.

Unveiling mechanisms underlying short- and long-term memory (STM and LTM) and consolidation of STM into LTM is an important issue in neuroscience. The role of cAMP-PKA-CREB pathway in consolidation of Pavlovian conditioning is conserved among Aplysia, flies and mammals. Our results may provide a foundation for further genetic approach to reveal the mechanisms underlying consolidation. It was known since the original report of thermotaxis that temperature preference reaches a local maximum 2–3 hours after cultivation temperature is shifted and that the preference is then once slightly weakened and finally reaches a plateau (Hedgecock and Russell, 1975, Figure 1E). *slo-2(nj131gf)* animals were slow to change behavior whereas *cng-3(jh113)* animals could transiently acquire an ability to perform thermotaxis right after cultivation temperature was shifted but gradually lost the ability as cultivated longer at the new temperature. One possibility is that two different components, one being fast but transient and the other late but persistent, might exist to shape the transition of temperature preference (Figure S7D). Our results suggested that SLO and CNG channels suppress the fast component and that the late component is dependent on CNG-3 but not on SLO channels since *slo-1; slo-2* double if mutants showed normal thermotaxis when cultivated for a long time at new temperature. Revealing mechanisms by which each component is regulated, for instance by a suppressor screen for *cng-3* mutants or analysis of animals in a transient state of consolidation, might lead to comprehensive understanding of consolidation of STM into LTM.

Vertebrate CNG channels, which are essential for functions of sensory neurons such as rod and cone photo receptors and olfactory receptor neurons, are also expressed in the central nervous system and play versatile roles (Podda and Grassi, 2014). CNG channels with different subunit composition have different channel properties such as sensitivity for cNMP and ion permeability, and function in different cell types (Kaupp and Seifert, 2002). Our and others’ results provide clues to understand how CNG channels with distinct subunit composition contribute to different cellular processes. Among CNG channels in *C. elegans*, TAX-2 and TAX-4 are essential for thermotaxis as well as for chemotaxis (Coburn and Bargmann, 1996; Komatsu et al., 1996), while CNG-3 seems to play a rather auxiliary role (O’Halloran et al., 2017). Although an *α* subunit TAX-4 can form a functional homo-tetramer that is far more sensitive to cGMP than the heteromer consisting of TAX-4 and a β subunit TAX-2 (Komatsu et al., 1999), TAX-2 that cannot form functional homomer by itself is still essential for thermosensation and chemosensation, indicating that heteromer-specific functions exist. CNG-3 was suggested to form a hetero-tetramer with TAX-4 and TAX-2 in AWC (O’Halloran et al., 2017), implying that a similar heteromer also forms in AFD. Considering our results suggesting that CNG-3 is necessary for steady state but not for transient state of thermotaxis, investigating how different composition of CNG channels function during such specific behavioral or cellular contexts in combination with analysis of their electrophysiological property would provide further understanding on roles of CNG channels in functions of the nervous system.

Our results suggested a possibility that the preference transition in thermotaxis might be useful as a model system for early onset epilepsies caused by Slack/*KCNT1* mutations. Given that quinidine, a blocker of Slack channel (Bhattacharjee et al., 2003; Yang et al., 2006), was not effective for every case of Slack/*KCNT1*-related epilepsies (Bearden et al., 2014; Chong et al., 2016; Mikati et al., 2015), and that quinidine also blocks other ion channels and may cause cardiac toxicity (Yang et al., 2006), screening for drugs that act more specifically on Slack using *C. elegans* would be valuable. Since Slack is expressed in cardiomyocytes (Yuan et al., 2003), identification of molecules that functionally interact with Slack only in neurons using suppressor screen against *slo-2(nj131gf)* animals would provide better drug targets.

This study particularly focusing on a transient state of decision making not only provided scientific insights into mechanisms that determine the timing of decision making, but also revealed a possibility for a clinical application that may contribute to the treatment of early onset epilepsies.

## Author Contributions

I.A. designed and performed experiments, wrote the manuscript, and secured funding.

M.T., T.S. and K.I. performed experiments.

Y.K. and S.N. provided expertise and feedback.

I.M. supervised and secured funding.

## Acknowledgements

We thank T. Tamada for technical assistance on experiments; H. Matsuyama for performing Fourier transformx; K. Noma and Y. You for critical reading and intensive discussion; A. Hart for valuable suggestions. Some strains were provided by the *Caenorhabditis* Genetics Center (CGC), which is funded by NIH Office of Research Infrastructure Programs (P40 OD010440). This work was supported by JSPS KAKENHI Grant Numbers JP 26870266, JP 22123010, JP 16H01272 and JP 16H02516.

## STAR Methods

### CONTACT FOR REAGENT AND RESOURCE SHARING

Further information and requests for resources and reagents should be directed to and will be fulfilled by the Lead Contact, Ikue Mori.

### EXPERIMENTAL MODEL AND SUBJECT DETAILS

*C. elegans* strains were cultivated on nematode growth medium (NGM) plates seeded with *E. Coli* OP50 strain (Caenorhabditis Genetics Center (CGC), Twin Cities, MN, USA) as described (Brenner, 1974). N2 (Bristol) was used as the wild type strain unless otherwise indicated. Transgenic lines were generated by injecting plasmid DNA or PCR fragments directly into hermaphrodite gonad as described (Mello et al., 1991).

### METHOD DETAILS

#### Behavioral assays

Population thermotaxis (TTX) assays were performed as described previously (Ito et al., 2006). Briefly, animals cultivated at 17°C or 23°C or subjected to a shift of cultivation temperature were placed on the center of the assay plates without food with the temperature gradient of 17–23°C and were allowed to freely move for 60 min. The assay plate was divided into eight sections along the temperature gradient, and the number of the adult animals in each section was scored. Ratio of animal numbers in each section was plotted in histograms. Thermotaxis indices were calculated as shown below:
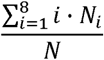

N_i_: number of animals in each section i (i = 1 to 8), N: total number of animals on the test plate.

#### Forward genetic screen for mutant animals slow to change thermotaxis behavior

For mutagenesis, wild type animals were treated with 50 mM ethyl methanesulfonate (EMS, Nacalai, Kyoto, Japan) for 4 hours at room temperature. F1 generation of the mutagenized animals were cultivated at 17°C for 5 days and were allowed to self-fertilize and give rise to F2 progeny. F2 animals were then transferred to 23°C, cultivated for 3 hours and then put on a thermal gradient for thermotaxis assays. While most of the animals migrated to the warm region of the thermal gradient, animals that still migrated to the cold region, which were putatively slow to change behavior, were isolated. To discriminate animals that are slow to change behavior from those that constitutively prefer cold temperature, strains isolated above were cultivated constantly at 23°C and were subjected to thermotaxis assays. Animals that migrated to the cold region when cultivated at 17°C and then at 23°C for 3 hours but migrated to the warm region when cultivated constantly at 23°C were considered to be slow to change behavior.

#### Suppressor screen against *slo-2(nj131gf)*

*slo-2(nj131gf)* animals were mutagenized with EMS as described above. Mutagenized animals were cultivated at 17°C for 5 days then at 23°C for 3 hours and subjected to thermotaxis assays as described above. While most of the animals still migrated to the cold region of the thermal gradient, animals that migrated to the warm region as if wild type were isolated.

#### Mapping of *nj131* and *nj172*

We crossed *nj131* animals with the wild type polymorphic CB4858 strain (Hillier et al., 2008). Since the Slw phenotype of *nj131* was inherited semi-dominantly, we isolated F2 animals that showed either Slw or wild type phenotype and identified crossover sites essentially as described (Wicks et al., 2001). We mapped *nj131* to a 1.3-Mb interval between nucleotides 11, 136, 567 and 12, 419, 537 on linkage group (LG) *X*.

For the mapping of *nj172*, we first crossed *slo-2(nj131gf)* animals four times with CB4858 to obtain a strain in which *slo-2(nj131gf)* allele exists on a background of CB4858 genome (IK2092). We then crossed *nj172; slo-2(nj131gf)* with IK2092, isolated F2 animals in which Slw phenotype was suppressed and identified crossover sites. We mapped *nj172* to a 1.1-Mb interval between nucleotides 8, 723, 437 and 9, 847, 895 on LG *IV*.

#### Whole-genome sequencing

Genomic DNA was purified with Puregene core kit A (Qiagen, Hilden, Germany). Genome was sequenced with MiSeq (Illumina, San Diego, CA, USA), and the sequence was analyzed with CLC Genomics Workbench (CLC bio, Aarhus, Denmark).

#### Plasmids

pOX nSlo2 was a gift from Larry Salkoff (Addgene plasmid #16202) (Yuan et al., 2000). A DNA clone including a part of *cng-3* cDNA (yk348a9) was provided by Yuji Kohara. To generate plasmids to express *slo-2* or *cng-3* cell-specifically, we fused promoter sequences of *gcy-8, ceh-36, ttx-3, lin-11, glr-3 or myo-3*, cDNA of *slo-2* or *cng-3* and the *unc-54* 3'UTR sequence by MultiSite Gateway Technology (Thermo Fisher Scientific, Waltham, MA, USA). To generate plasmids to express SLO-2 in HEK293T cells, *slo-2* cDNA was inserted in pCAG vector. Details of the plasmid constructs are available from the authors.

*Peft-3::Cas9-SV40 NLS::tbb-2 3'-UTR* (Addgene plasmid #46168) and *PU6::unc-119_sgRNA* (#46169) were gifts from John Calarco (Friedland et al., 2013). *unc-119* sgRNA sequence in *PU6::unc-119_sgRNA* was replaced with sequences of sgRNAs for *unc-22, slo-2* #1 and *slo-2* #4, which were GAACCCGTTGCCGAATACAC(AGG) (Kim et al., 2014b), GAAATACCTGGTTTTCGGGG(TGG) and GCACAATGCACGATGCAAGG(CGG), respectively.

pCFJ150 *pDESTttTi5605[R4-R3]* (Addgene plasmid #19329), pCFJ601 *Peft-3::Mos1 transposase* (#348874), pMA122 *Phsp::peel-1* (#348873), pCFJ90 *myo-2p::mCherry* (#19327), pCFJ104 *myo-3::mCherry* (#19328) and pGH8 *rab-3p::mCherry* (#19359) were gifts from Erik Jorgensen (Frøkjær-Jensen et al., 2008, 2012). A plasmid containing *slo-1* cDNA was a gift from Jonathan T. Pierce (Davis et al., 2014). *gcy-8* promoter sequence, *slo-1* cDNA and *unc-54* 3'UTR sequence were fused into pCFJ150 to make pIA109,

#### Knockout of *slo-2* by CRISPR/Cas9

*slo-2* gene was knocked out by CRISPR/Cas9 essentially as described (Farboud and Meyer, 2015; Kim et al., 2014b). *slo-2(nj131gf)* animals were injected with *Peft-3::Cas9-SV40 NLS::tbb-2 3'-UTR*, pRF4 *rol-6(su1006, gf)*, pSN680 *unc-22* sgRNA, pIA009 *slo-2* sgRNA #1 and pIA054 *slo-2* sgRNA #4. *unc-22/+* and *unc-22/unc-22* F1 twitcher animals were subjected to genotyping for *slo-2* gene locus. Although deletion between the sites of sgRNA #1 and #4 was not detected, small InDels around the site of sgRNA #1 were detected by polyacrylamide gel electrophoresis (PAGE) following PCR that amplified 60 bp region including the site of sgRNA#l with primers TCGTGATTTCTTGGAAGAATTCT and tggtgtgttactgaaatacCTGG. A strain that homozygously carried a small deletion at the *slo-2* locus was isolated from progenies of a *unc-22/+* animal with a small deletion. Sequencing of genomic *slo-2* locus revealed that this strain has a deletion of 2 nucleotides in the seventh exon, which causes a frame shift after His344. No InDel was found adjacent to the site of sgRNA #4. *unc-22* mutation was then kicked out by picking non-twitcher animals.

#### Single copy insertion by MosSCI

Single copy of *gcy-8p::slo-1a(+)* was inserted to a genome essentially as described (Frøkjær-Jensen et al., 2008). *ttTi5605;unc-119(ed3)* animals derived from EG6699 strain (Frøkjær-Jensen et al., 2012), which show Uncoordinated (Unc) locomotion, were injected with a mixture of pCFJ601, pIA109, pMA122, pGH8, pCFJ90 and pCFJ104. NGM plates with non-Unc animals were heat-shocked at 42°C for 2 hours. Non-Unc animals without mCherry fluorescence were selected and subjected to genotyping. PCR was performed to confirm an insertion of a full-length transgene.

#### Imaging analyses

Calcium imaging was performed as described elsewhere (Kobayashi et al., 2016; Yoshida et al., 2016). A single adult animal that expressed genetically encoded calcium indicators R-CaMP2 (Inoue et al., 2014) in AFD and GCaMP3 (Tian et al., 2009) in AIY or an animal that expressed GCaMP3 and tagRFP in AFD was placed on a 10% agar pad on a cover slip with 0.1 μM polystyrene beads (Polysciences, Warrington, PA, USA) and covered by another cover slip for immobilization (Kim et al., 2013). The immobilized animals were placed on a Peltier-based temperature controller (Tokai Hit, Fujinomiya, Japan) on a stage of BX61WI microscope (Olympus, Tokyo, Japan). The red and green fluorescence was separated by the Dual-View optics system (Molecular Devices, Sunnyvale, CA, USA), and the images were captured by an EM-CCD camera C9100-13 ImageEM (Hamamatsu Photonics, Japan) at 1 Hz frame rate. Excitation pulses were generated by SPECTRA light engine (Lumencor, Beaverton, OR, USA). The fluorescence intensities were measured by the MetaMorph imaging system (Molecular Devices).

Expression of SLO-2::mCherry was observed with BX53 upright microscope (Olympus).

#### Electrophysiology

HEK293T cells were transfected with plasmids encoding yellow fluorescent protein (YFP) and either wild type or H159Y mutant form of SLO-2b with lipofectamine2000 (Invitrogen). The macroscopic current was recorded from HEK293T cells expressing YFP by whole cell patch clamp configuration using Axopatch 200B amplifiers, Digidata 1322A and the pClamp 9 software (Axon Instruments, Foster City, CA, USA) at room temperature, as previously described (Tateyama and Kubo, 2013). Bath solution contained (in mM) 140 NaCl, 4 KCL, 10 HEPES, 1 CaCl_2_, 0.3 MgCl_2_ (pH= 7.4). Pipette resistances ranged from, 1 to 3 MQ when the glass pipette was filled with the pipette solution that contained (in mM) 130 KCl, 10 NaCl, 10 HEPES, 5 K_2_ATP, 4 MgCl_2_, 3 EGTA, 0.3 GTP and various concentration of CaCl_2_ (pH= 7.3). To achieve0, 0.2, 0.6, 2, 6 and 20 μM of free [Ca^2+^]_i_, 1, 1.96, 2.55, 2.86, 2.99, 3.10 mM of CaCl_2_ was added to the pipette solution, according to Ca-Mg-ATP-EGTA Calculator vl.O using constants from NIST database #46 v8 (http://maxchelator.stanford.edu/CaMeATPEGTA-NIST.htm). After the rapture of the sealed membrane, cells were held at −60 mV for at least 3 min to dialyze the pipette solution into the cell, and then depolarizing step pulses (10 mV increment for 200 ms) were applied. Immediately after the step pulse applications, cells were held at 0 mV (50 ms) for the tail current analysis. The amplitudes of the currents were measured at the end of step pulses and normalized by the membrane capacitance in each cell. The current densities were then averaged and plotted as a function of the membrane potentials. To analyze the voltage dependence of the channels, the tail current amplitudes at the very beginning of 0 mV were measured and normalized by the maximal one at the most depolarized potential (G/G_max_). G/G_max_ values were plotted against the membrane potential and fitted to a Boltzmann function to estimate the membrane potential of a half maximal activation (V_1/2_) in each cell.

#### RT-PCR

mRNA was extracted from *C. elegans* by RNAiso PLUS reagent (Takara Bio, Kusatsu, Japan) and then subjected to reverse transcription by ReverTra Ace^®^ qPCR RT Master Mix with gDNA Remover (Toyobo, Osaka, Japan). cDNA was then subjected to real time PCR with THUNDERBIRD^®^ SYBR qPCR Mix (Toyobo) on CFX96 Real-Time PCR Detection System (Bio-rad, Hercules, CA, USA). Sequences of primers used for qPCR were listed in Supplemental Table 1. Quantities of housekeeping genes such as *act-1, gpd-1 and lmn-1*, which encode actin, GAPDH and lamine, respectively, were used as internal controls.

### QUANTIFICATION AND STATISTICAL ANALYSIS

Error bars in histograms and line charts indicate standard error of mean (SEM).

In boxplots, the bottom and top of boxes are the first and third quartiles, and the band inside the box is the median. The ends of lower and upper whiskers represent the lowest datum still within 1.5 interquartile range (IQR), which is equal to the difference between the third and the first quartiles, of the lower quartile, and the highest datum still within 1.5 IQR of the upper quartile. For multiple comparison tests, one-way analyses of variance (ANOVAs) were performed, followed by tests either with Tukey-Kramer or by Dunnette methods as indicated in each figure legend. Welch Two Sample t-test was used to compare two values.

*References in Key Resources Table and supplemental figure legends

(Consortium, 2012)

(Davies et al., 2003)

(Wang et al., 2013)

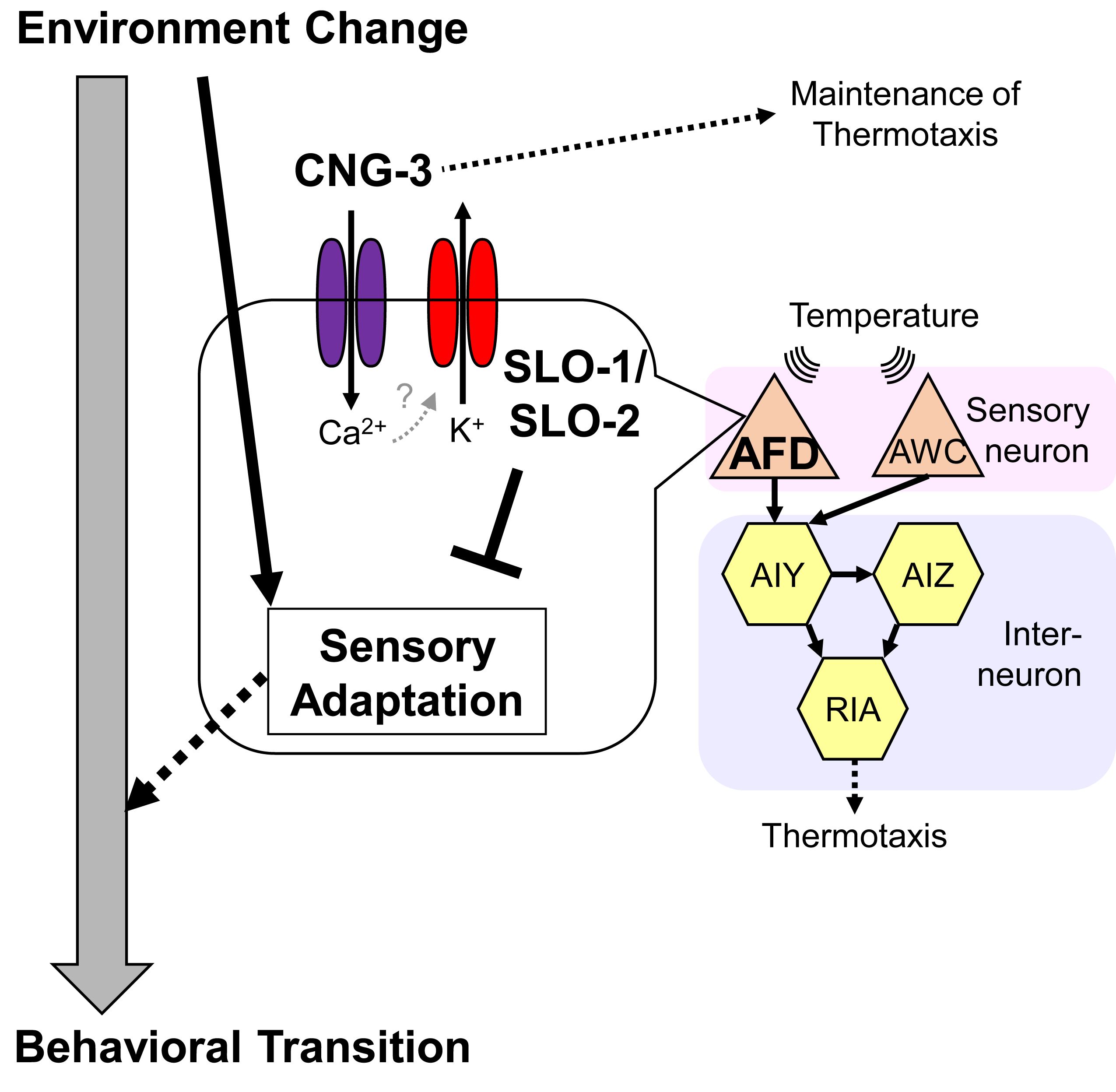

